# On simulating cold stunned turtle strandings on Cape Cod

**DOI:** 10.1101/418335

**Authors:** Liu Xiaojian, James Manning, Robert Prescott, Zou Huimin, Mark Faherty

**Affiliations:** School of Remote Sensing and Information Engineering, Wuhan University, Wuhan, China; National Oceanic Atmospheric Administrations’s Northeast Fisheries Science Center, Woods Hole, Massachusetts, United States of America; Massachusetts Audubon’s Wellfleet Bay Wildlife Sanctuary, Wellfleet, Massachusetts 02663 United States of America; Shandong Agricultural University, Taian, China

**Keywords:** Kemp’s ridley turtles, FVCOM model, ocean current, particle tracking

## Abstract

Kemp’s ridley turtles were on the verge of extinction in the 1960s. While they have slowly recovered, they are still endangered. In the last few years, the number of strandings on Cape Cod Massachusetts beaches has increased by nearly an order of magnitude relative to preceding decades. This study uses a combination of ocean observations and a well-respected ocean model to investigate the causes and transport of cold-stunned animals in Cape Cod Bay. After validating the model using satellite-tracked drifters and local temperature moorings, ocean currents were examined in the Cape Cod Bay in an attempt to explain stranding locations as observed by volunteers and, for some years, backtracking was conducted to examine the potential source regions. The general finding, as expected, is that sub 10.5°C water temperatures in combination with persistent strong wind stress (>0.4Pa) will result in increased strandings along particular sections of the coast dependent on the wind direction. However, it is still uncertain where in the water column the majority of cold stunned turtles reside and, if many of them are on the surface, considerable more work will need to be done to incorporate the direct effects of wind and waves on the advective processes.

## Introduction

Kemp’s ridley is the smallest and the most endangered sea turtle in the world and shows little sign of recovery [1,2,3]. Their perilous situation is attributed primarily to the over-harvesting of their eggs during the last century. In 1970, it was listed by the U.S. Fish and Wildlife Service [4] as “endangered throughout its range” and it has received Federal protection under the U.S. Endangered Species Act of 1973 ever since [2].

The adult Kemp’s ridley turtles are mainly active in coastal waters less than 165 feet deep primarily in the Gulf of Mexico where satellite telemetry is used to understand their migratory patterns and habitat for the past few decades [5,6,7,8,9]. The smaller juvenile Kemp’s ridley are found in shallow waters, often foraging in less than 3 feet of water [10]. Juveniles are also found in the Gulf of Mexico [11,12] as well as along the eastern seaboard of the United States as far north as Nova Scotia, Canada [13] leaving warm seas to feed on crabs and other prey[14].

Sea turtles on the East Coast of the US swim to warmer waters to spend the winter. Hart et al [15] discuss the seasonal variability of both Loggerhead and Kemp’s ridley strandings based on drifter bottles and particle tracking. As in the loggerhead study by Santos et al [16], the objective is often to document the location and probable cause of mortality [17] using a variety of methods such as deployment of coded oranges [18]. In many cases, it is abrupt changes in temperature that trigger mortality. Turtles in the vicinity of Virginia, tend to migrate south in late fall each year [19,20] and, even in Florida, significant mortalities have been reported [21] as a result of being caught in some cold water bodies/estuaries.

In laboratory experiments, Schwartz [22] demonstrated that Kemp’s ridleys become inactive at temperatures below 10°C, although they are somewhat more resistant than other species. If a sea turtle becomes cold stunned there is little chance of survival without assistance. Morreale, et al. [23] studied cold stunned stranding events i n the late 1980s on Long Island, NY. Burke et al [24] specifically addressed the causes of these events and found one year, in particular, when fewer strandings occurred. They attributed this inter-annual variability to slight changes in the prevalent wind direction at the critical time of year when the water temperature fell below 10°C.

When Kemp’s ridley’s inside Cape Cod Bay in Massachusetts are late in heading south and the water temperature gets below 10.4°, they become “cold stunned” [25]. They are apparently trapped due to the presence of the Cape Cod landmass. These turtles, typically 2 to 3 years old, are sized at 26.9 mean straight carapace length [25] between a dinner plate and a serving platter [26]. They are so small satellite telemetry devices can not be easily used to track the turtles like that larger individuals in Gulf of Mexico [11,12,27]. In recent years, a larger number have been stranding on the shores of Cape Cod. During the summer months there are a few strandings reported because of turtles getting a) entangled in fishing gear, b) hit by a boat, and c) stuck in sandbars on a falling tide but these are less important later in the year when the majority of the turtles strand due to the cooler temperatures. These turtles are extremely weak, often wash ashore, and need to be rescued by dozens of trained volunteers each fall. Since 1979, Wellfleet Bay Wildlife Sanctuary staff and volunteers have patrolled the beaches of Cape Cod, on the lookout for cold-stunned turtles, which are rapidly transported to the New England Aquarium for evaluation and rehabilitation. In the last few years, the number of strandings has increased by nearly an order of magnitude relative to the strandings in the preceding decades.

This paper explores the transport of cold stunned Kemp’s ridleys using the Finite Volume Community Ocean Model (FVCOM), a numerical ocean model well-suited to simulating coastal ocean processes [28]. After validating FVCOM by using drifter data and mooring data (see supplementary material), we look at the number of sea turtles stranded in various towns along the shore of Cape Cod Bay from 2012 to 2015 and try to explain these distributions based on FVCOM-derived simulated flow fields, particle tracks, wind records, and observations of water temperature.

## Methods

### FVCOM model

FVCOM is a prognostic, unstructured-grid, Finite-Volume, free-surface, three-dimensional (3-D) primitive equations Community Ocean Model developed originally by Chen et al [1,28]. As validated in the supplementary material, the ability of FVCOM to accurately solve scalar conservation equations in addition to the topological flexibility provided by unstructured meshes and the simplicity of the coding structure has made FVCOM ideally suited for many coastal and interdisciplinary scientific applications [29].

FVCOM model has been run on several grids spanning the Northeast Continental Shelf of the US and is still evolving. The third generation grid (GOM3) was used in this experiment covering Gulf of Maine and Georges Bank waters and hourly flow fields are available from 1978 to 2015. The GOM3 grid was developed primarily for the prediction of coastal ocean dynamics, with horizontal grid resolution ranging from 20 meters to 15 kilometers and includes tides. The vertical domain was divided into 45 s-coordinate levels. We also experimented on finer-resolution “MassBay” grid that had hourly flow fields available from 2011-2014.

### Drifter observations

The surface drifters used in this study track the top meter of the water column [30]. This type is typically referred to as the “Davis-style” or “CODE” drifter first developed for the Coastal Ocean Dynamics Experiment in the early 1980s (Fig 1) [31]. Our version of this drifter typically has a metal-frame and four fabric sails measuring 91 cm long and 48 cm wide. These student-built units were deployed primarily by local fishermen in the vicinity of Cape Cod from 1978 to 2010 with tracks archived for public access. More than a thousand of these drifters have been deployed off the New England coast in the past few decades [32,33]. The primary purpose of these drifters is to validate ocean models (see supplementary material).

**Fig 1.**
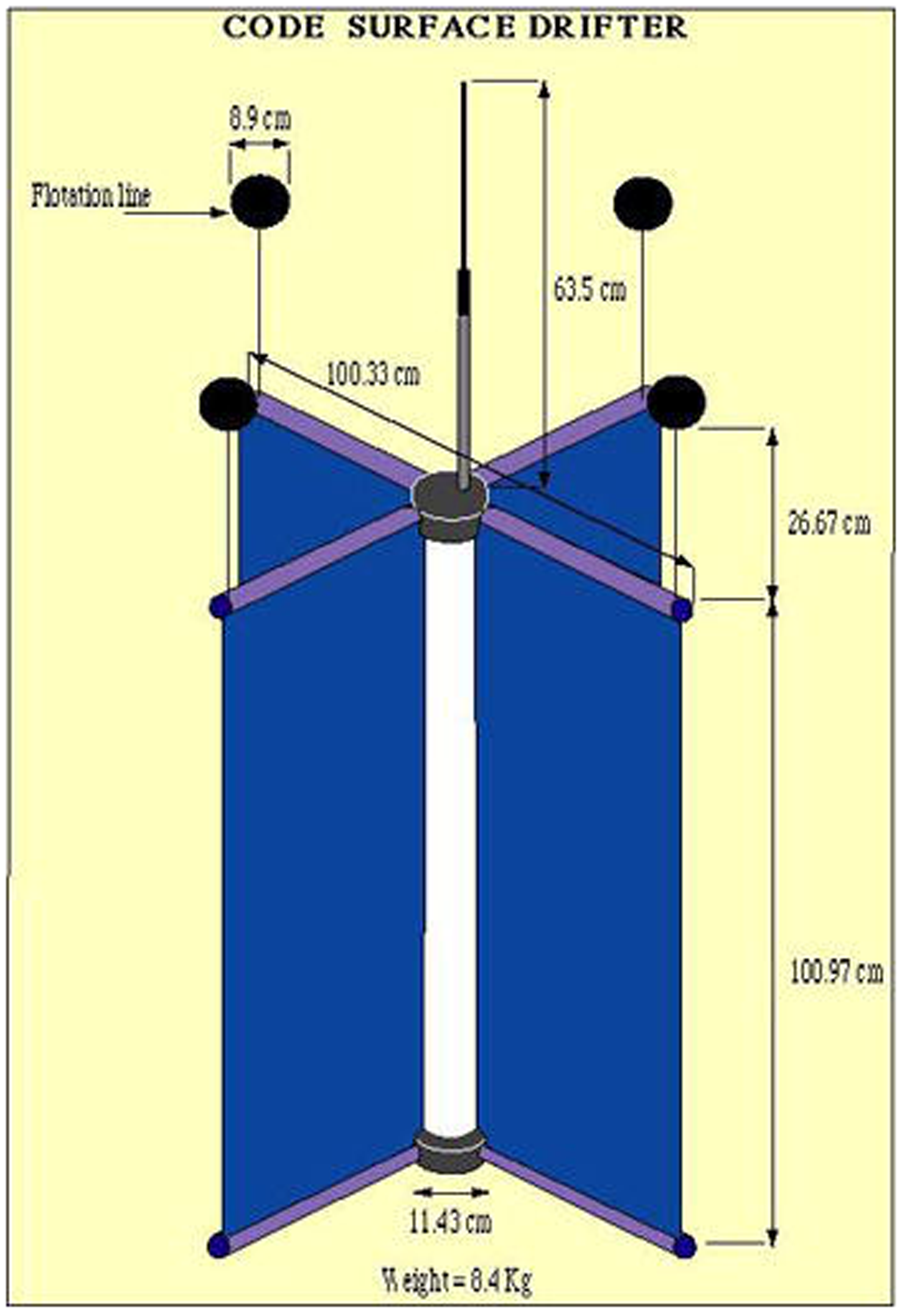
Schematic of standard surface drifter from Poulain [30] (left) and photo of surface drifter used in this report being recovered after stranding in Barnstable MA (right).

### Moored temperature observations

There are at least three different sets of moored ocean temperature observations available in the Northeast US. The Northeastern Regional Association of Coastal Ocean Observing System (NERACOOS) typically has several mooring sites in the Gulf of Maine [34]. In 2017, they added a mooring (CDIP) inside Cape Cod Bay. The NERACOOS Mooring A location is shown in Fig 2. The Environmental Monitors on Lobster Trap Project (eMOLT) [35] comprises another collection of moorings with 18+ years of hourly bottom temperatures from dozens of locations off the New England coast including several within Cape Cod Bay. Finally, the Mass Division of Marine Fisheries (DMF) maintains several diver-serviced bottom temperature time series at a set of shipwrecks off the Massachusetts coast.

**Fig 2.**
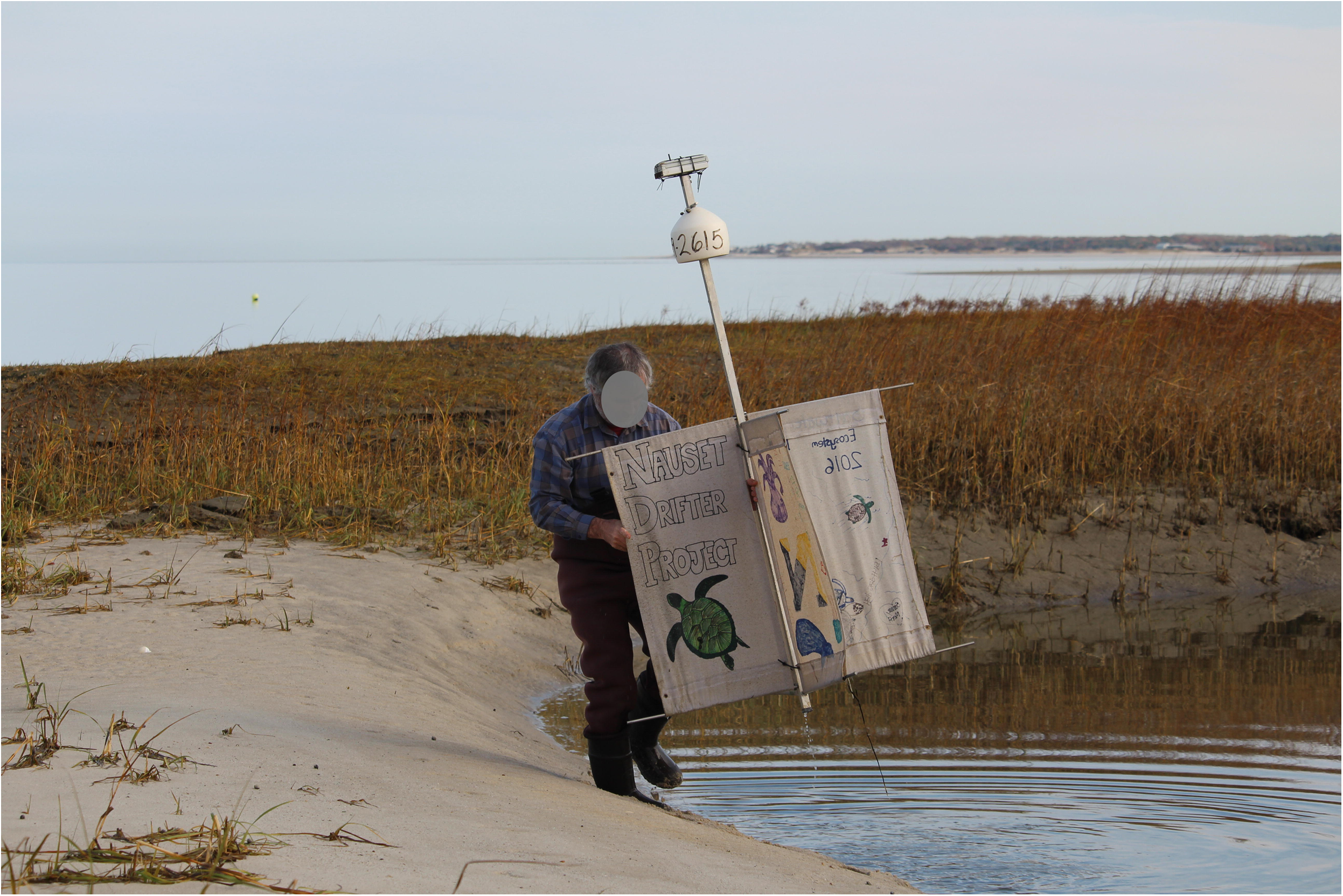
Locations of moored observations used in this report including two NERACOOS moorings (blue triangles), five eMOLT moorings (red circles) and four Massachusetts Division of Marine of Fisheries moorings (green squares). Also posted are the positions where wind and surface temperature was extracted from the models.

### Simulating forward and backward turtle trajectories

The science of tracking particles through the coastal ocean is still evolving [29] and there are a variety of techniques [25,33,34] that can be applied depending on the physical and biological behavior of the particle/animal under investigation. In this case, since the animal’s position in the water column and its ability to regulate that depth is unknown, we present here a very simple case of moving it through the model’s surface fields using the nearest grid nodes with hourly time steps with no consideration of other factors such as wind and waves. The comparison of observed and modeled particle tracks is presented in supplementary material Appendix I.

For the case of backtracking the cold stunned turtles to estimate their origin, we first need to get the simulated turtle off the beach in order to begin moving it backward through the water. The process of selecting a “short distance off the beach” is not easy. We experimented with multiple methods (simple to complex) to solve this problem (see supplementary materials). For the purpose of this study, we chose the simplest method which involves a) finding the nearest point on the model-based coastline to the stranding location, b) calculating the line from the specified stranding to that point, and c) then extending the line ***X*** km out to sea. After experimenting with several values, we chose to set “***X*** =1.5 km”. As suggested in Nero et al [27], the value selected is partly a function of the bathymetric slope offshore where for their case “X” varied from 0.3km to 1.0km. The backtracking is terminated when the estimated water temperature exceeds the cold-stunned level. For reasons discussed below, we chose this to be 10.5 °C. We backtracked turtles for 2012 and 2013^1^. We selected the 167 turtle strandings in these years who’s simulated trajectory stayed off the beach. In other words, we focus on only those 167 cases where the backtracking generated origins within the bay.

### Binned averaged current and wind

In order to examine the surface current conditions in the bay for a specified period of time, we bin-averaged both observed and modeled flow fields in on 0.05 ° bins. The “observed current” is derived from averaging the first differences of the near-hourly drifter fixes within each bin. The model data is the hourly output of the FVCOM model as described in section 2.1 above. We averaged the observed current, modeled current, and the model wind stress. The “modeled current” in each grid cell is the average velocities of all nodes within each bin whenever drifters were present in the bin. While technically different types of current velocity (Lagrangian vs Eulerian), we assume they are the same. After removing the tidal effects by sequentially averaging over the solar (24 hour) and lunar (24.841) periods, we calculated the average surface current of each bin. The wind stress was derived from the same weather model that drives the FVCOM ocean model which is local implementation of the NCAR Mesoscale Model (MM5).

### Turtle stranding data

The Kemp’s ridley turtle data used in this experiment was provided by Massachusett Audubon’s Wellfleet Bay Wildlife Sanctuary. These data include information on date and location of stranding for each cold-stunned Kemp’s ridley sea turtle from 1991 to 2015. Hundreds of trained volunteers have been participating in this project since 1979. Beach walkers cover the coastline each fall to document strandings and report their findings. In order to quantify the relationship between wind stress and the number of strandings, we calculated the correlation between wind stress and the number of strandings when the temperature was below 10.5°C. As noted above, it is not only the strength of the wind that may be important but the persistence of the wind over time. For this reason, we chose to calculate the “3-day sum of the windstress components”.

### Sea-surface temperature observations

In addition to the modeled sea-surface temperature (SST) estimates available hourly, we also have observations of SST from satellite. While clear images are only available a few times per month in this region, they provide another snapshot realization of the conditions in Cape Cod Bay. For the purpose of this report, we accessed the raw data prepared by the University of Delaware and posted via the MARACOOS.org website.

## Results

### Kemp’s ridley strandings on Cape Cod

Fig 3 shows the number of Kemp’s ridleys stranded on the shores of Cape Cod Bay from 1991 to 2015 (with some years 1994-1998 not available due to lack of staff during those times). As shown, the number of strandings vary from year to year but have increased dramatically in recent years.

**Fig 3.**
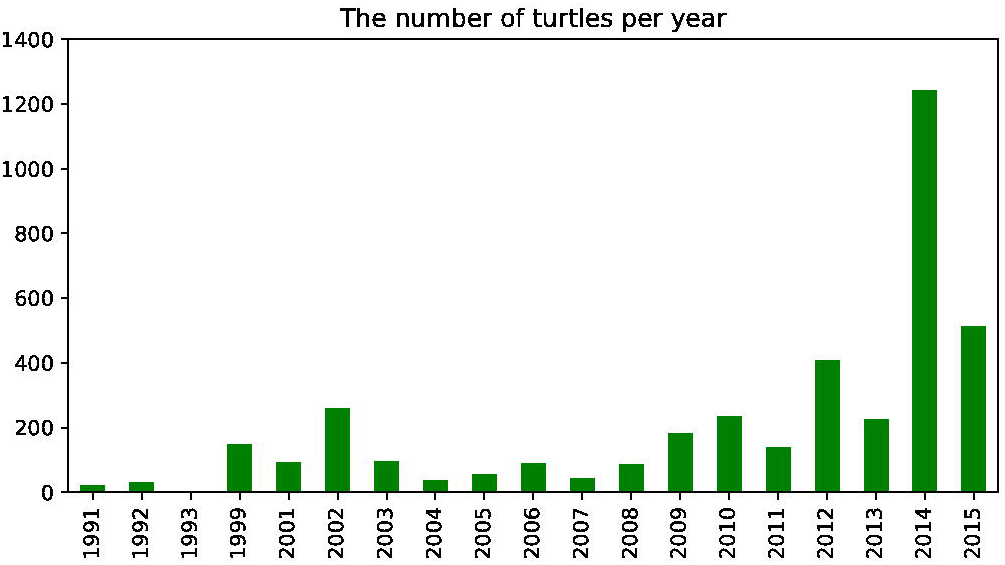
1991-2015 Cape Cod Bay turtle strandings.

Turtles were recovered in several towns each year between 2012 and 2015 (Figs 4 and 5). The towns with the largest numbers of strandings between 2012-2015 were located on the mid to outer Cape. Among them, each town reported strandings in more than one year of the study period, and strandings were reported every year in Wellfleet and Brewster. While most stranding events occur on the Outer Cape, some occur Mid Cape. Our hypothesis is that year-to-year changes of stranding locations are primarily due to small changes in the surface current.

**Fig 4.**
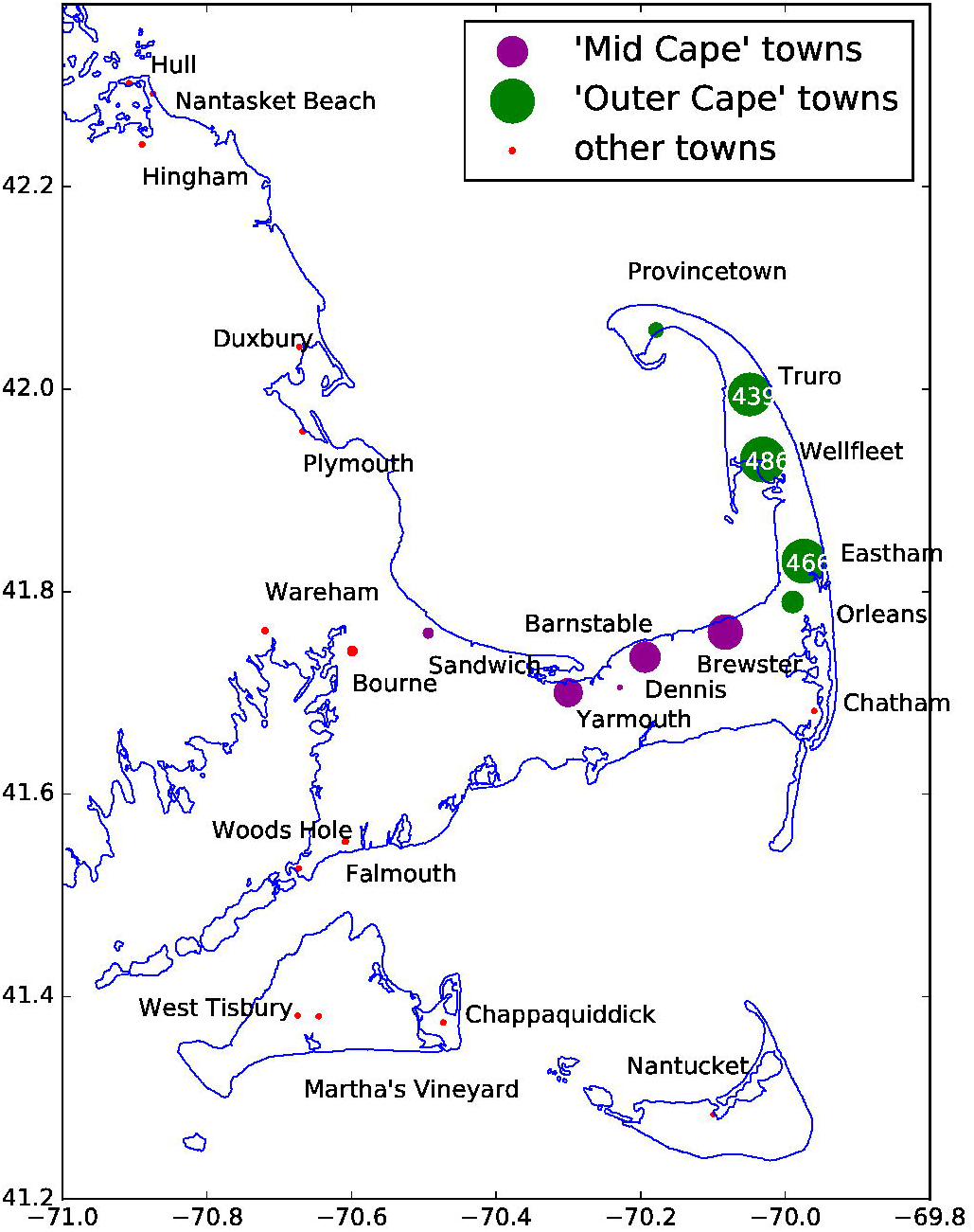
Total turtle strandings per town from 2012-2015. Most of the “Outer Cape” towns (green dots) had >400 strandings while some of the “Mid Cape” towns (purple) had ∼200.

**Fig 5.**
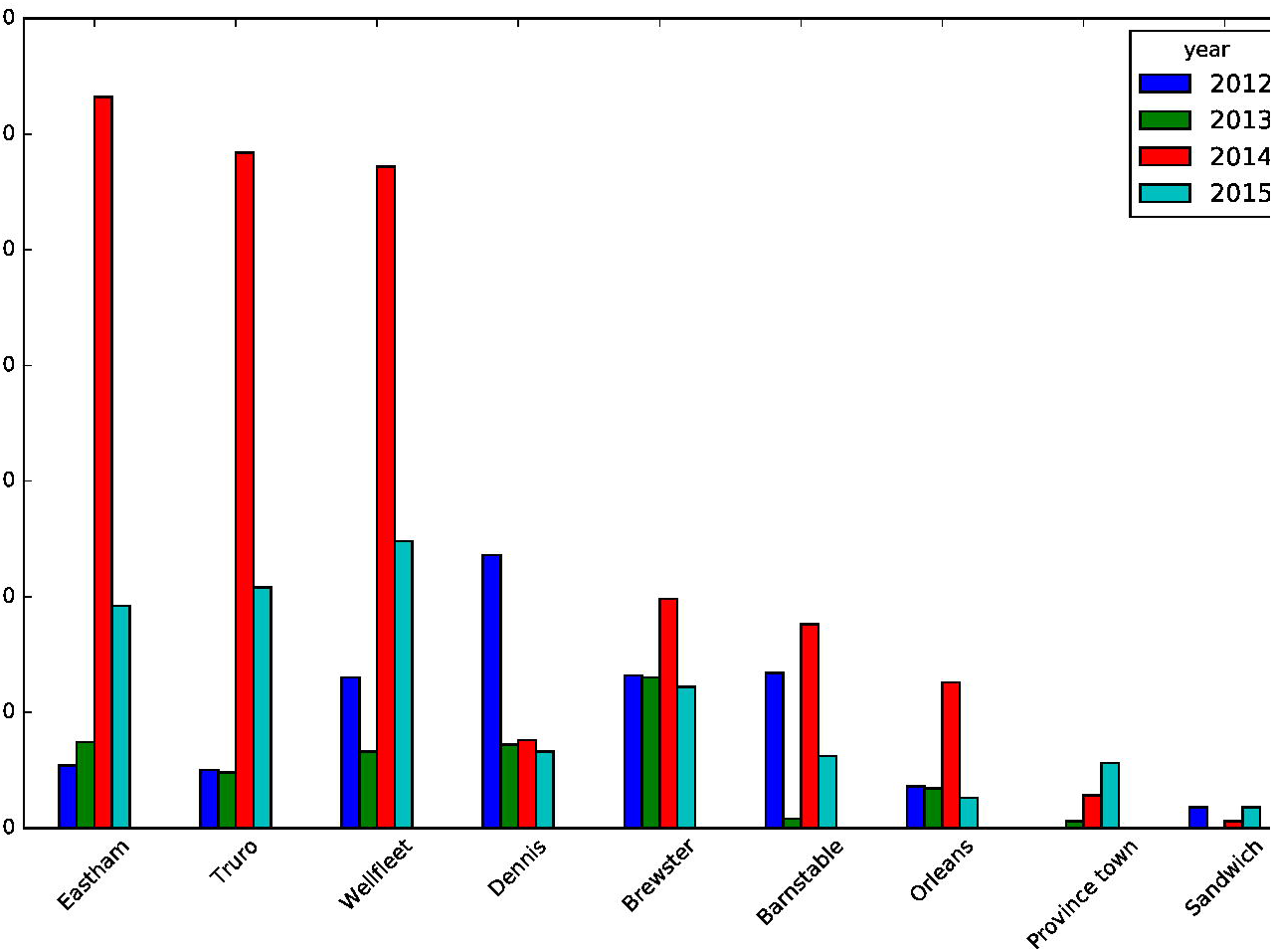
Chart of the number of strandings by town between 2012 - 2015 showing most in the first three Outer Cape towns.

### Examining binned current and wind

Despite recent efforts to survey the bay from both ships and planes, we do not know the exact location of the cold stunned turtles in the bay or in the water column. However, considering that the cold-stunned turtles are not capable of swimming, we assume they generally follow the surface currents with some effects by the wind and waves. It is expected that more skillful predictions of stranding positions may be made using observed ocean surface currents and wind stress to numerically advect water parcels forward. Fig 6 shows the number of turtles stranded on the shores of Cape Cod Bay per day in late 2014 with most occurring the week of November 21st. Fig 7 maps the mean current and the mean wind stress from November 18 – 23, 2014, and the distribution of the stranded sea turtles for these six days.

**Fig 6.**
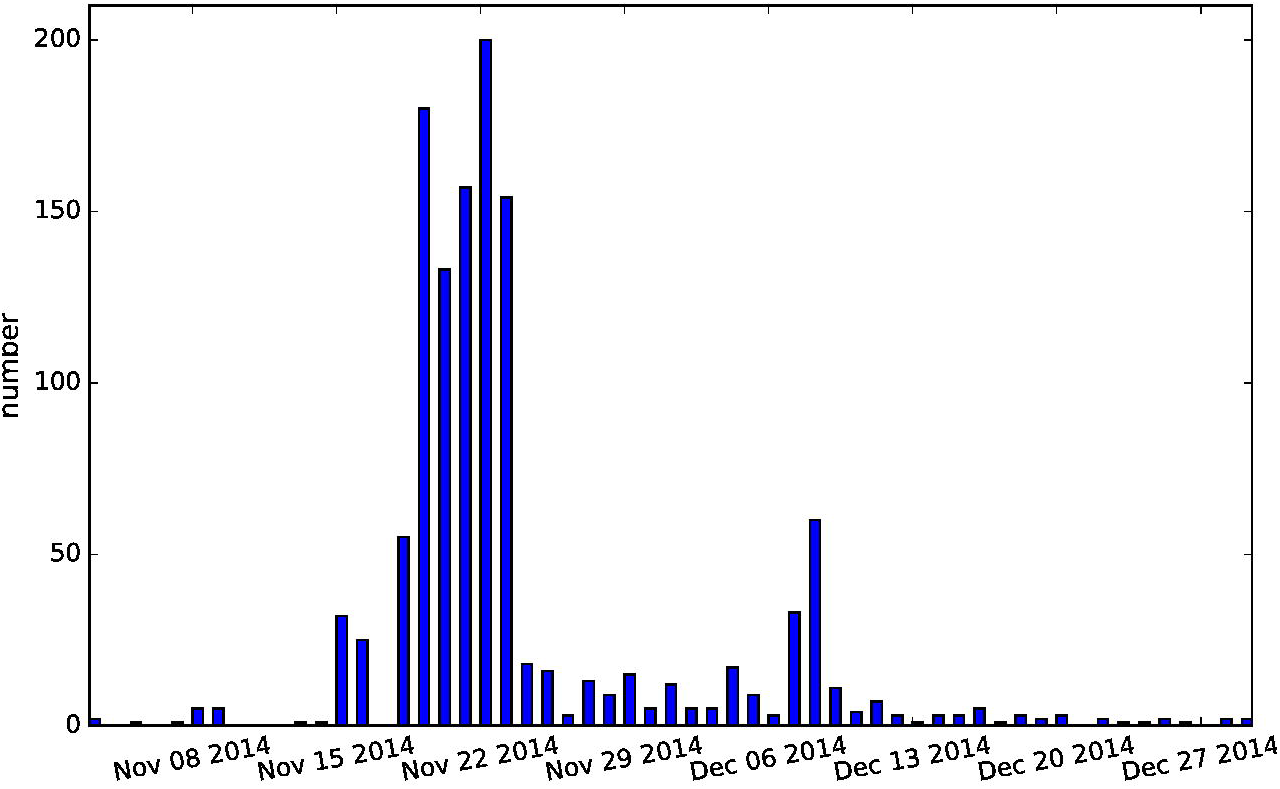
The number of turtles stranded on Cape Cod Bay beaches in Nov-Dec 2014.

**Fig 7.**
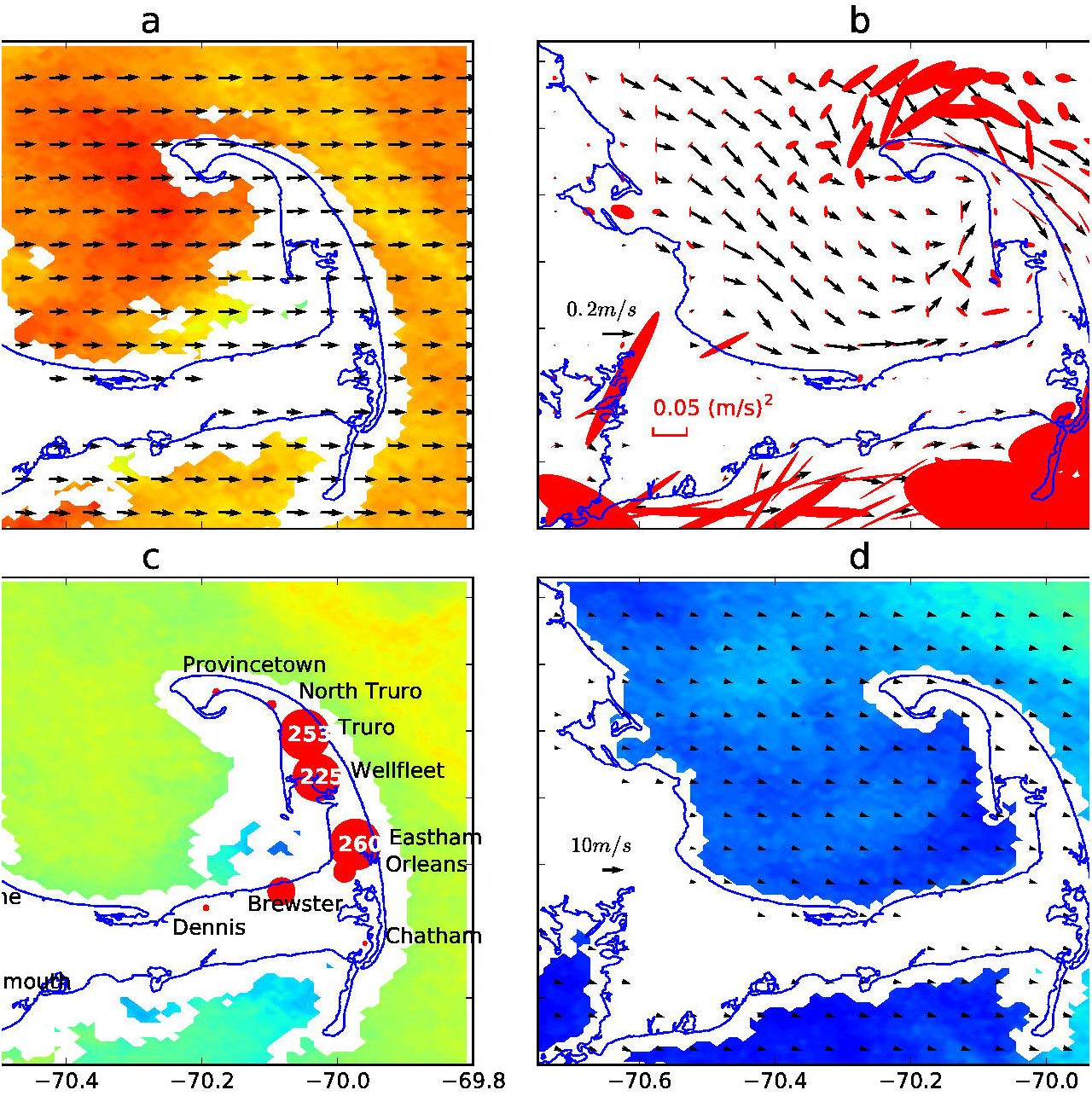
November 18-23, 2014 mean wind (black arrows) with November 3, 2014 sea surface temperature (a), Mass Bay grid simulation of mean ocean surface currents (black arrows) during that time overlaid on surface current variance ellipses (pink) (b), and distribution of turtle strandings by town with November 19, 2014 sea surface temperature (c), and mean wind for the entire month of November 2014 with December 26, 2014 sea surface temperature (d). The legends are posted on the land mast on the western side of the charts. The largest red dot in panel c denotes 260 turtles stranded in Eastham during the period 18-23 November 2014.

From November 18, 2014 to November 23, 2014, as depicted on the turtle stranding map (Fig 7c), Truro, Wellfleet, and Eastham had the most strandings (253, 225, and 260, respectively), and their positions are consistent with the oceanic currents and winds shown by the simulated current and wind maps (Fig 7a and 7b), distributed in the area where currents and wind stress are most intense, and in their direction. It can be seen that the combination of ocean current and wind calculated by FVCOM model can be used to explain the general distribution of strandings. Comparing the mean wind for one week in mid-November 2014 (Fig 7a) to the mean wind for the entire month of November 2014 (Fig 7d), we see the intensity of the mid-November period explains the unusually high rate of strandings on the Outer Cape. The intensity of these events is best described in the form of time series plots below. In the future turtle rescue work, FVCOM 3 day forecast model can be used to simulate stranding events and help volunteer beach walkers focus their efforts.

As previous studies have found, when the water temperature of the Cape Cod Bay is less than 50F (10℃), the metabolism of Kemp’s ridley turtle will slow to the point where they are unable to move and are at the mercy of the wind, waves, and currents [25]. As shown in Fig 2, we set up 10 temperature detection points and 10 wind stress monitoring points across Cape Cod Bay and Mass Bay to extract the temperature and wind stress simulated by the model. In the following plots (Figs 8–10), we examine the relationship between wind stress and strandings especially when the ocean temperature is below 10.5 °C.

**Fig 8.**
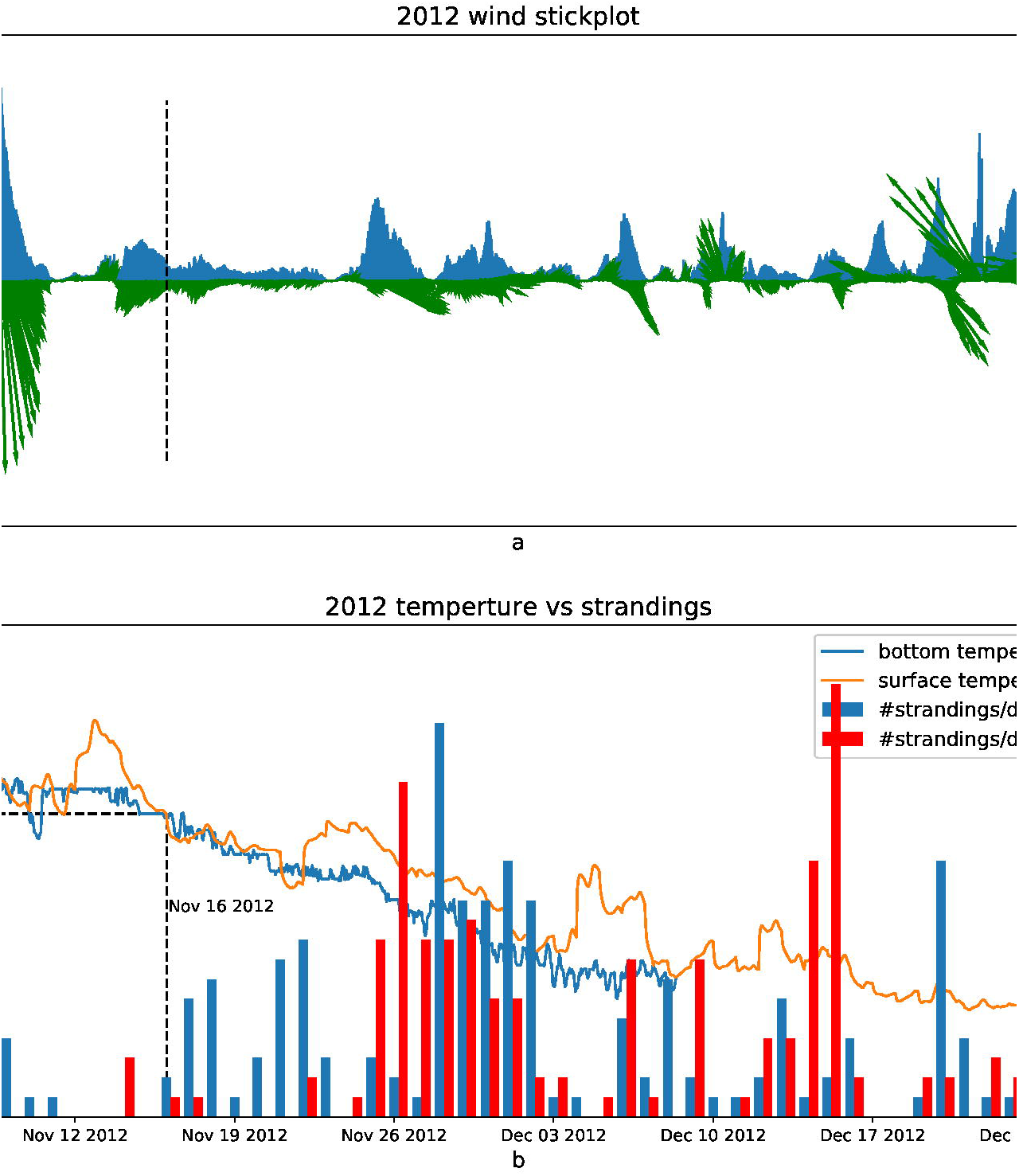
2012: Top panel shows wind stress (Pa) direction shown in green vectors (north up) and magnitude in purple. Bottom panel shows Mid-Cape strandings (blue bars) the Outer-Cape strandings (red bars), modeled surface temperature (green line), and observed bottom temperature (blueline) in 2012.

**Fig 9.**
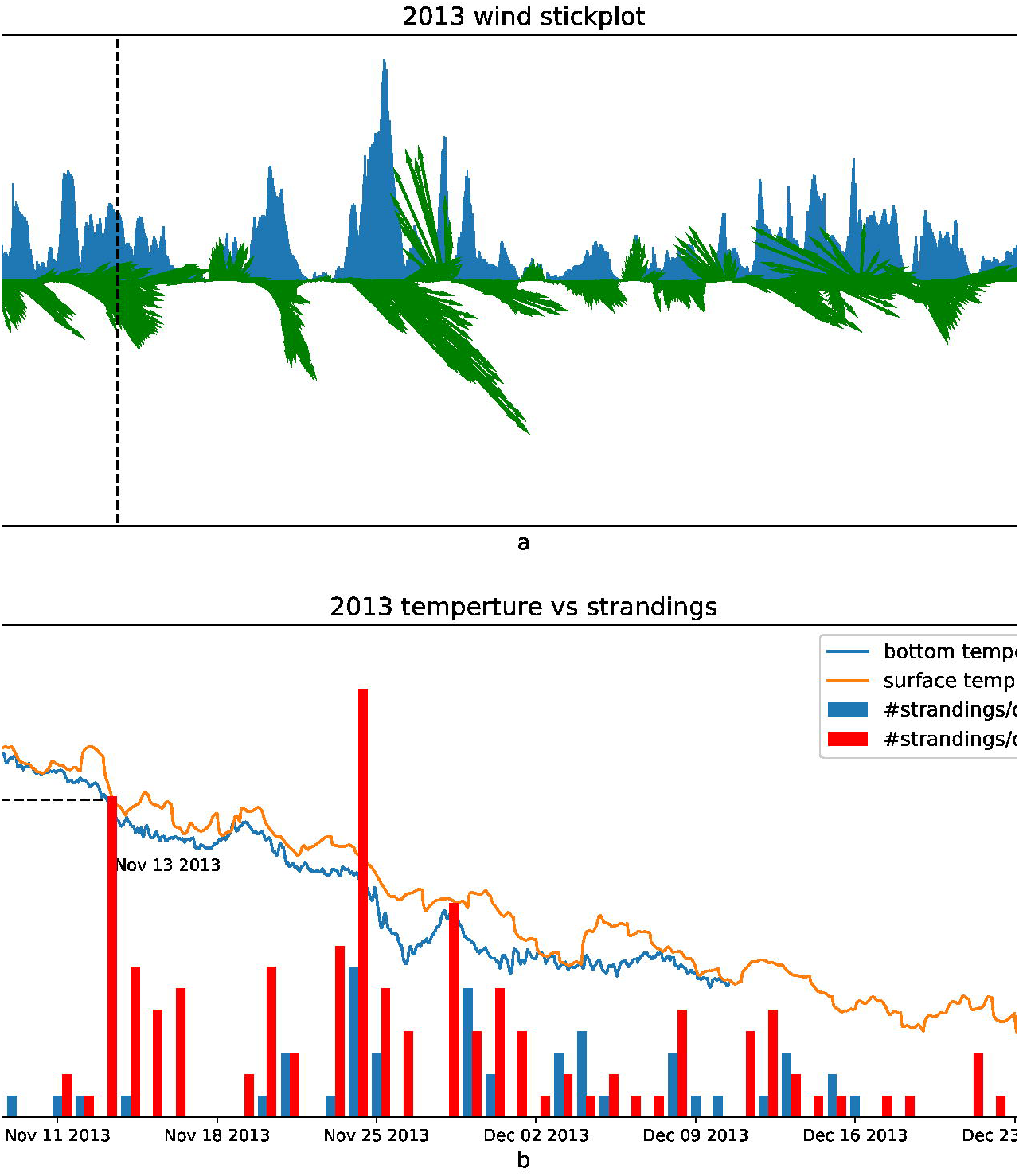
2013: Top panel: Wind stress with direction shown in green vectors (north up) and magnitude in purple. Bottom panel: Mid-Cape (blue bars) and Outer-Cape (red bars) strandings, modeled surface temperature (green line), and observed bottom temperature (blueline) in 2013.

**Fig 10.**
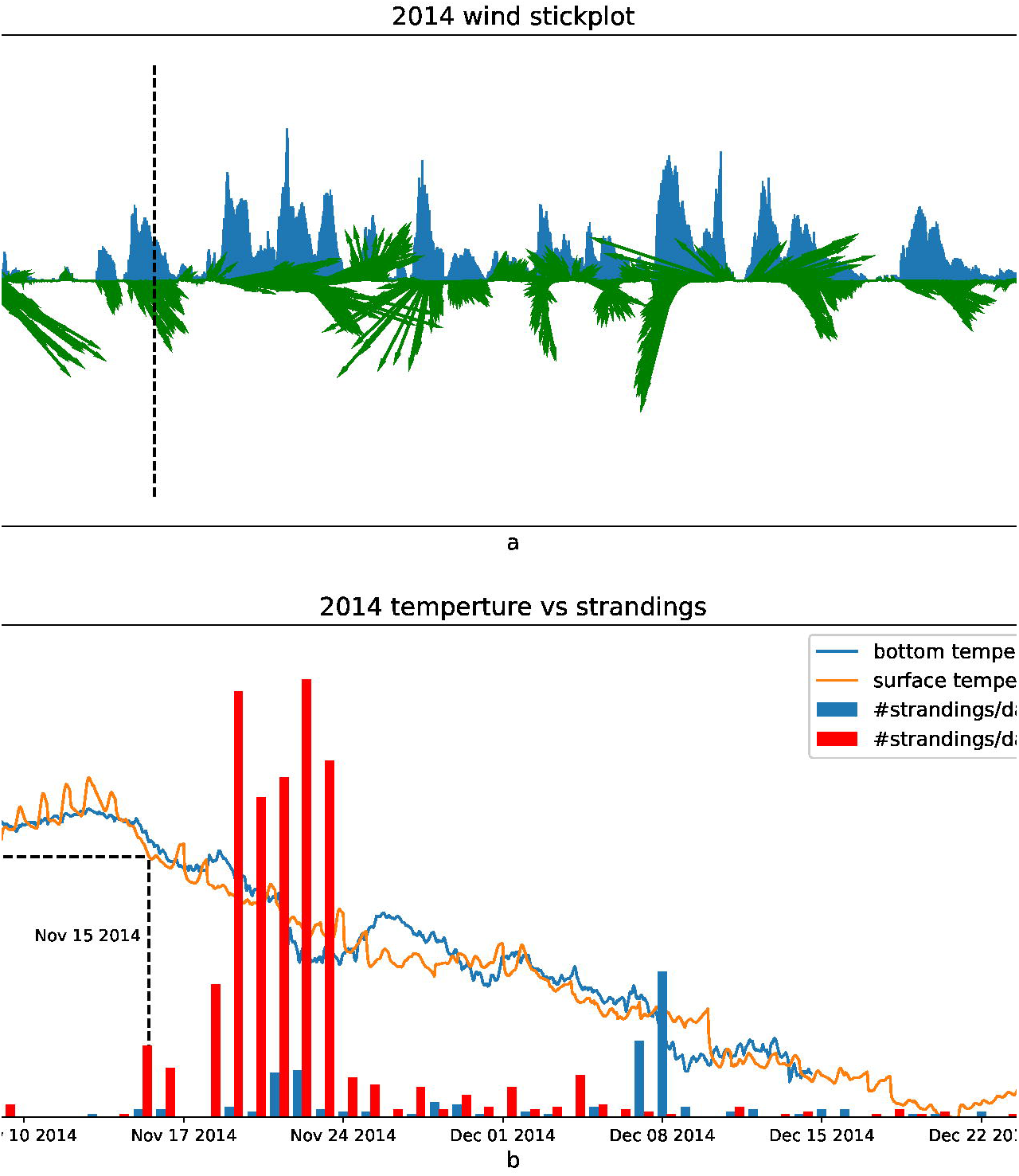
2014: Top panel: Wind stress with direction shown in green vectors (north up) and magnitude in purple. Bottom panel: Mid-Cape strandings (blue bars), Outer-Cape strandings (red bars), modeled surface temperature (green line), and observed bottom temperature (blueline) in 2014.

Figs 8–10 show the relationship between the wind stress (top panels), and the number of strandings in “Outer Cape” and “Mid-Cape” (bottom panels) towns in 2012, 2013, and 2014, respectively. The average of the surface temperature simulated by all the FVCOM model points (specific model sites shown in Fig 2) and the average of the bottom temperature observed in Cape Cod Bay (specific observation sites shown in Fig 2) are also plotted in the bottom panels. The wind stress vector is the average simulated by FVCOM model points (sites shown in Fig 2). When the temperature is less than 10.5 °C and the wind stress from a particular direction is greater than ∼0.4 Pa, the strandings generally increase but both conditions must be met. In the case of 2012 (Fig 8), for example, the strong wind in early November resulted in near-zero strandings since the water temperature was still above 11°C. In the case of 2013 (Fig 9), there was a pair of events with strong northwesterly wind at the end of November that coincided with a drop in temperatures resulting in peaks in strandings around 26 November and 1 December. Note the strong winds (>0.4Pa) in mid-December 2013 resulted in very few strandings presumably due to the turtles having fled or already stranded by that time. In the case of 2014 (Fig 10), the wind in mid-to-late November was not strong but it was persistently from the west over many days with a drop in both surface and bottom temperature of a few degrees which evidently resulted in historically high amounts of strandings.

Fig 11 (left hand panels) shows the correlation between the number of turtle strandings in “Outer Cape” towns and the 3-day sum of the **eastward** wind stress in 2012, 2013, 2014 and mean respectively. Fig 11 (right hand panels) show the correlation between the number of turtles strandings in “Mid Cape” towns and the **southward** wind stress in 2012, 2013, 2014, and mean respectively. In most cases, the correlation is greater than 90% and exceeds 95% in several cases. Wind stress and the number of strandings per three days show a linear correlation.

**Fig 11.**
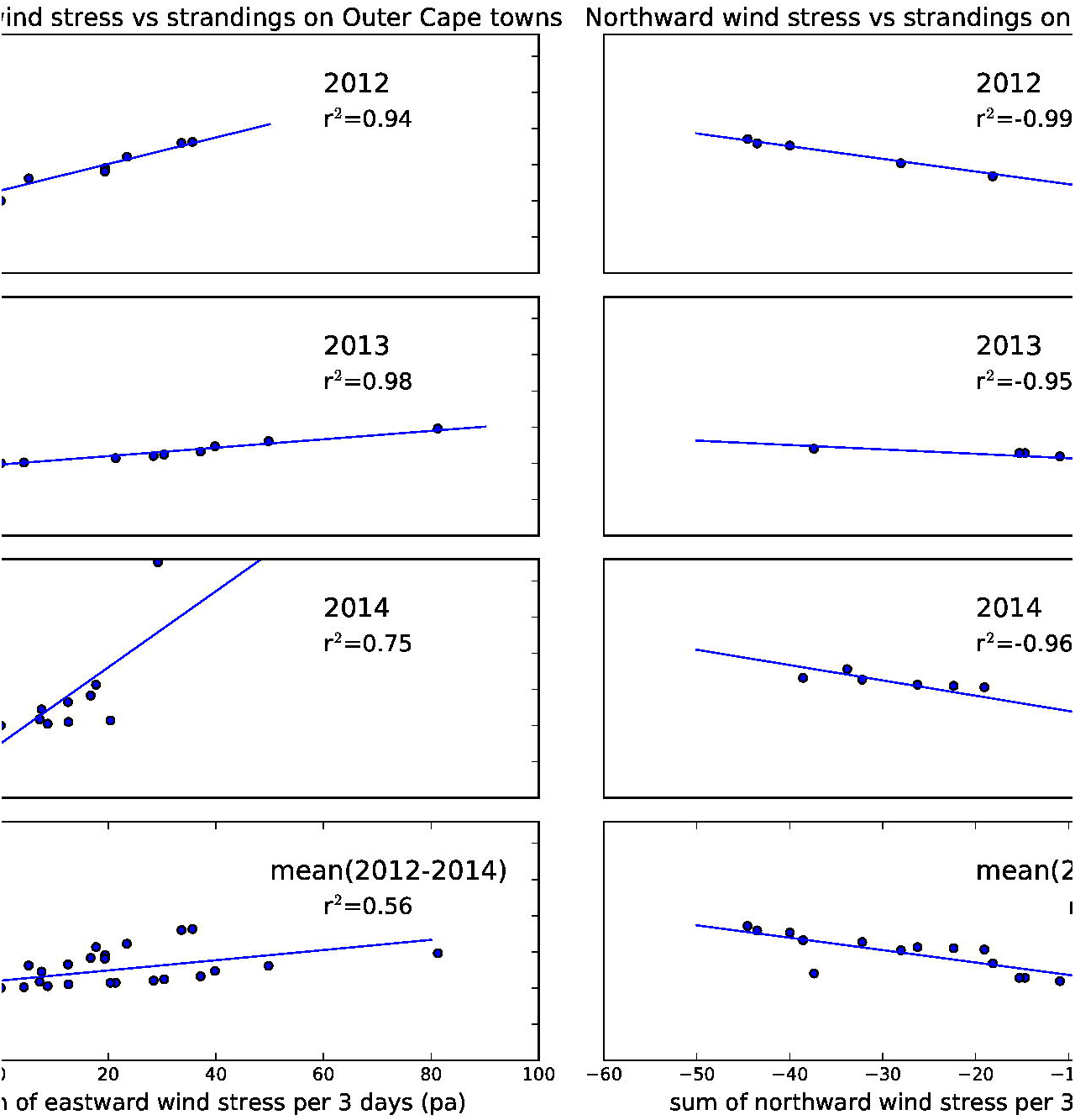
Linear correlations between eastward (left) and southward (right) components of the wind stress and number of strandings (summed over 3 days) in 2012, 2013, 2014 and mean (2012-2014) for Outer Cape (left) and Mid Cape (right) towns.

Finally, in order to estimate the source of turtles (ie their distribution/origin at the time of first being cold stunned), we look at the backtracking results (Fig 12). We can also look at the estimate of temperature along their track (Fig 13). In these time series we see that a) many of the stranded turtles were exposed to <10.5 °C water in mid-to-late November and b) there were more December stranding events in 2012.

**Fig 12.**
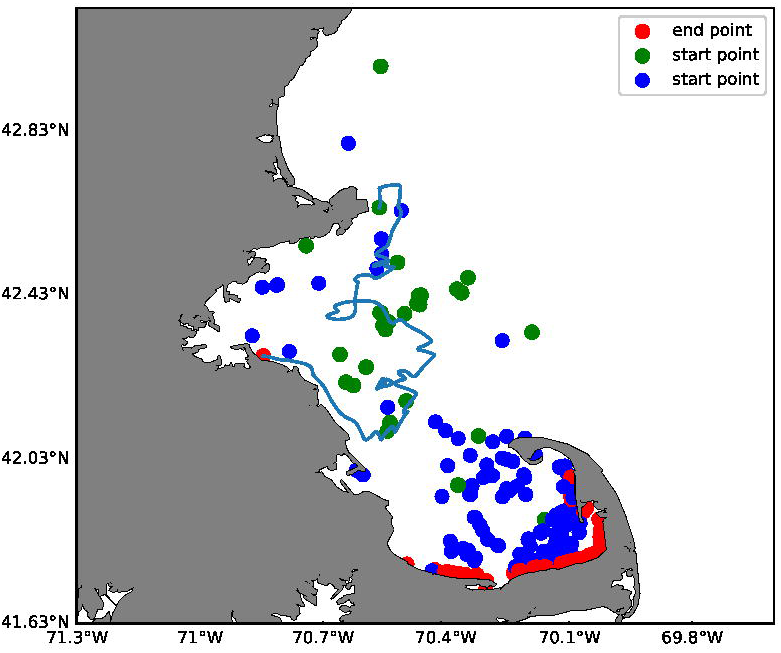
The backtracked positions of the turtles in 2012 and 2013. Red dots show landing locations in November and December. Blue and green dots show estimated locations of back-tracked turtles that stranded in November and December, respectively, when they first encountered <10.5°C. water. This assumes passive (non-swimming) turtles were only transported by the current when the modeled surface temperature is less than 10.5 °C. The blue line is one example of backtracking a turtle that landed just south of Boston Harbor in December. According to the model, that turtle was first cold-stunned off Gloucester, MA.

**Fig 13.**
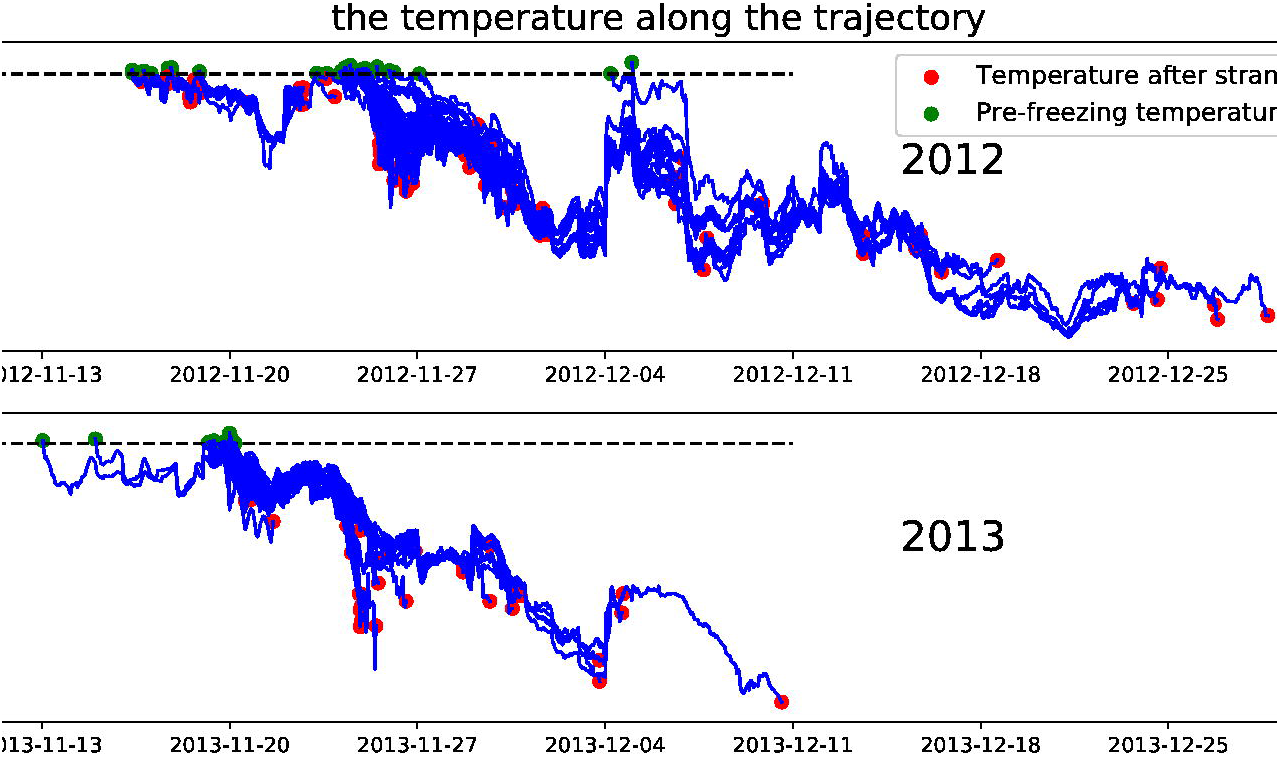
Modelled temperature of backtracked surface particles in 2012 (top panel) and 2013 (bottom panel).

We might ask where were the turtles that stranded in November located when they were first exposed to <10.5 °C surface waters? Our results show that turtles stranded in November 2012 and 2013 may be cold stunned at a variety of locations (blue dots in Fig 12) primarily in Cape Cod Bay but some in Massachusetts Bay. Turtles stranded in December 2012 and 2013 (green dots in Fig 12) may originate even further away, some as far as northern Massachusetts Bay. After first being exposed to <10.5 °C surface waters, the turtles are evidently in temperatures that vary considerably but gradually decrease by a few degrees at most before stranding (Fig 13).

## Discussion

Although the state-of-the-art coastal ocean models like FVCOM can help simulate cold stunned turtle pathways, there are a variety of issues to consider: First, uncertainty in the numerically simulated drifter tracks result in differences of up to several kilometers per day relative to the observed satellite-tracked drifter tracks. Obviously, there is significant uncertainty in the backtracking results shown in Fig 12. The 200-meter-per-kilometer difference in modeled vs observed trajectory (as shown in the supplementary materials) grows over long distances and time so that the numerical circulation models and particle tracking routines will need to be improved considerably before these particle tracking tools can be used confidently and operationally. Simulated trajectories will also differ from actual trajectories due wind and waves influencing cold stun turtle trajectories. There is a growing body of literature on surface wave effects on the transport of surface particles. Much of this research comes from the coast guard search and rescue operations [36, 37] as well as recent studies on oil spill dispersion. Methods have been devised to add the effects of multiples forces including direct wind effect, breaking waves [38], Stokes drift [39], inertial effects [40], and these will need to be considered in future work. Since it is still uncertain where in the water column these cold stunned turtles reside, we chose to focus on surface current fields as a first approximation in the study but these results would change significantly for turtles floating directly on the surface.

Additional uncertainties are introduced with the tracking methodology near the boundary of the model grid (i.e. in shallow waters near the shoreline). We need to further explore algorithms for forward and backward particle tracking near the coast especially in this study area where there is a large expanse of wetting and drying tidal flats and close to 30cm tidal range. Improving our particle tracking models with both the effects of wind and waves and processes near the coast is the primary goal in the near future.

Despite these limitations, models like FVCOM are improving every year particularly with increasing grid resolution, incorporation of more realistic bathymetry, and more observations to assimilate. In order to examine the effect of different grid resolution, for example, we conducted a sensitivity study comparing particle tracks on the “GOM3” vs “MassBay” grid (Fig 14). It can be seen here that the particles released on the GOM3 grid stranded on the Cape Cod beaches faster than those released on the MassBay. While the final stranding locations would likely be similar, the particles on the latter grid were not yet stranded at the end of the week-long simulation.

**Fig 14.**
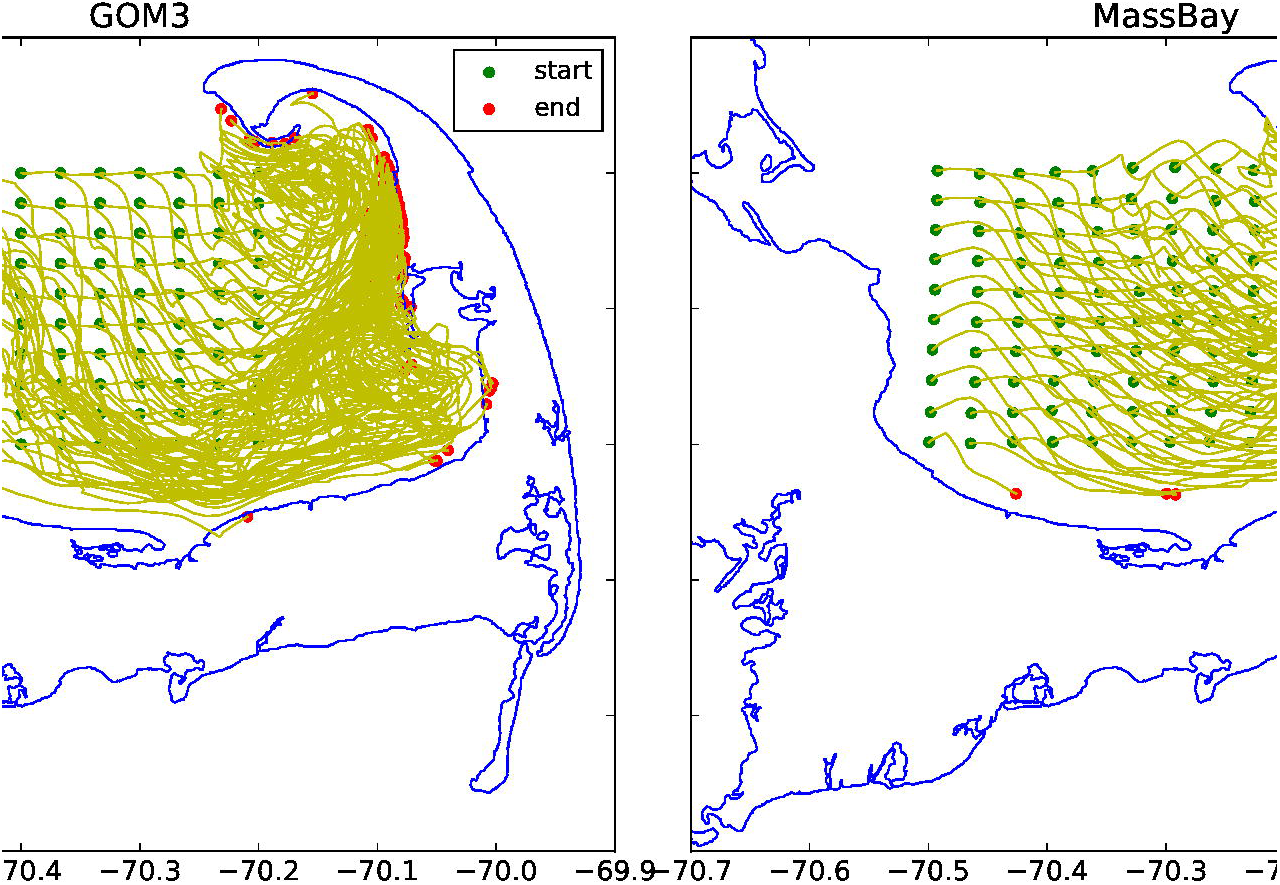
Comparing particle tracks on the GOM3 (left) and MassBay (right) grid during the week of 18-25 Nov 2014.

Still more observations are needed to monitor in-situ conditions throughout Cape Cod Bay including more realizations of the deeper currents not depicted by the standard surface drifters used here. There is uncertainty in both the deeper currents and the depth at which the turtles generally reside during this critical time in the fall. When we eventually know where these turtles originate and at what depth of the water column, divers or ROVs could be deployed to help locate the turtles. There is some chance that some turtles may be on the bottom waiting for the water to warm. Very few have been observed near the surface whenever searches have been conducted.

As the frequency of extreme weather events is apparently on the increase [41,42] the chances of turtles being suddenly surprised by wind and temperature changes will increase. As the population of animals continue to increase [3], as more individuals start their migration from the Gulf of Mexico beaches, and as more of these tropical species migrate north with warming conditions [43], will we see a larger sea turtle population in Cape Cod Bay in the future and will that result in larger strandings?

## Conclusion

With some specified degree of uncertainty, the ocean current and temperatures simulated by the FVCOM ocean model is similar to that observed by satellite-tracked drifters and moored observations. The model can help predict the strandings of the Kemp’s ridley turtle on the shores of Cape Cod and also be used to backtrack their source. In the forward prediction runs, the FVCOM model helped explain the anomalously large stranding event, for example, that occurred on Outer Cape Cod between November 18 – 23, 2014. In general, the geographic distribution of standings is consistent with the model simulation of ocean current and wind. Given these findings, coastal ocean models such as FVCOM can help explain the general locations of strandings and may help volunteers to rescue turtles in the future.

In the backtracking experiments, the FVCOM model was used to estimate the origin of stranded Kemp’s ridley turtles in 2012 and 2013, showing with a specified degree of uncertainty that most turtles stranded in November were originally located in a variety of locations in both Cape Cod Bay and Massachusetts Bay when they were first exposed to cold-stun temperatures (<10.5 °C). The turtles stranded in December may have originated well outside Cape Cod Bay just a few weeks earlier.

There are still many uncertainties as to when the turtles are cold-stunned but a simple regression of different components of the wind stress (with water temperatures below the 10.5°C threshold) vs number of strandings indicate that values greater than 0.4 Pa will induce more stranding events (r^2^=95). As expected, strong winds out of the west will result in Outer Cape strandings and strong winds out of the north will result in Mid-Cape strandings.

While progress has been made in simulating the stranding events, considerable more work needs to be conducted to a) determine the depth of the water column occupied by the turtles and b) the differential effects of surface advection due to wind and waves relative to near-surface currents.

## Supporting information

https://www.nefsc.noaa.gov/epd/ocean/MainPage/turtle/ccbay/S1_Supplementary_material_CCBay_manuscript.pdf

https://www.nefsc.noaa.gov/epd/ocean/MainPage/turtle/ccbay/S2_Supplementary_material_CCBay_manuscript.pdf

https://www.nefsc.noaa.gov/epd/ocean/MainPage/turtle/ccbay/S3_Supplementary_material_CCBay_manuscript.pdf

## Acknowledgements

Thanks to all those who have helped with this analysis, the many turtle rescue volunteers, the staff at Mass Audubon’s Wellfleet Bay Wildlife Sanctuary, Dr. Vitalli Sheremet for help in coding, anonymous reviewers at NEFSC, and Dr. Dai Honglei for long term encouragement and support. The Massachusetts Division of Marine Resources provided some of their moored temperature observations from Cape Cod Bay and the UMASS Dartmouth FVCOM modeling group provided on-line access to their simulations.

## Footnotes

1 Note: Since there were evidently issues with 2014 model hindcast, we chose to focus only on the previous two years in this analysis.

