## Supplementary material for "On simulating cold stunned turtle strandings on Cape Cod": https://www.nefsc.noaa.gov/epd/ocean/MainPage/turtle/ccbay/S1_Supplementary_material_CCBay_manuscript.pdf

### Appendix 1:

### More on circulation model and the validation of model results

#### More description of the numerical model

FVCOM is solved numerically by a second-order accurate discrete flux calculation in the integral form of the governing equations over an unstructured triangular grid. This approach combines the best features of finite-element methods (FE: grid flexibility) and finite-difference methods (FD: numerical efficiency and code simplicity). Relative to FE and FD, it provides a better numerical representation of both local and global momentum, mass, salt, heat, and tracer conservation (Chen and Beardsley, 2003). While grid size varies depending on the bathymetric complexity, the average for the grid used in Cape Cod Bay was 1.5 km.

#### Bin-averaged current and wind

We first examined the bin-averaged current and wind.

The current averaging is conducted for both the drifter observations and the model output and shown in Fig S1_1. The code to generate this figures and others in this Appendix is available at https://github.com/jamespatrickmanning/Lei_et_al_2019_CCBAY/tree/S1_model_vs_obs .

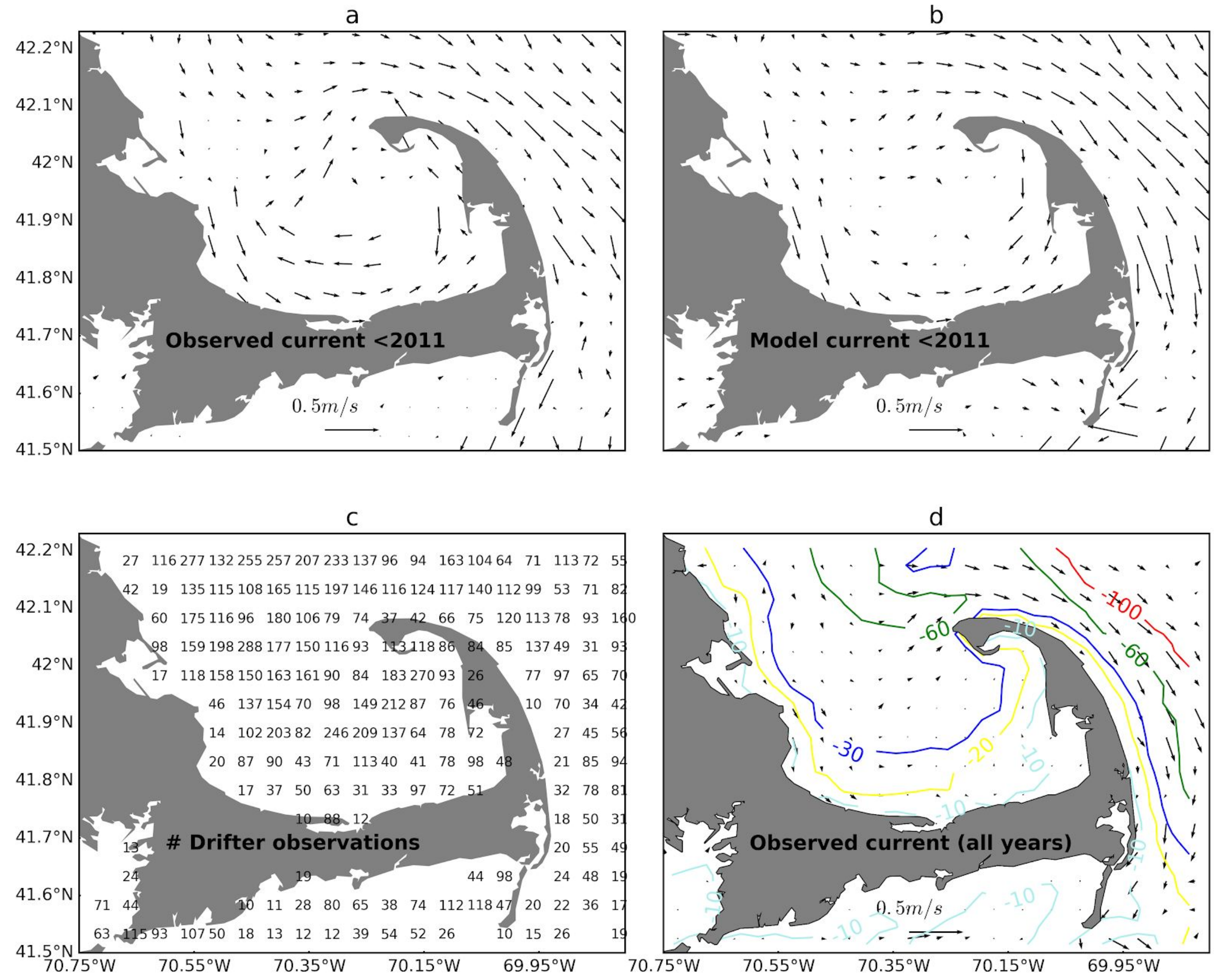


**Fig S1_1. Surface current derived from all drifters prior to 2011 (a) and FVCOM model-derived flow field at the same time as those drifter observations (b), number of drifter observations prior to 2018 (c), and mean field from all the drifters prior to 2018 with bathymetry contoured in meters (d).**

When we derive the mean field from drifters whenever there was GOM3 model data available (<2011), we get the vector plot Fig S1_1a. We can compare it to the model-derived field in Fig S1_1b. In a mean-flow sense, the model does well to estimate the flow of the coastal waters in this region. While there are some areas of discrepancies in, for example, the middle of the bay, most of the region is resolved by the model. When we calculate the overall mean differences (observed-mean), we find values of -0.02 and 0.03 m/s for the eastward and northward flows, respectively. As expected, the variability of the flow is less resolved by the model with the mean difference in standard deviations being 0.07m/s and 0.09m/s, respectively. Given additional drifter deployments in Cape Cod Bay providing more observations of current (Fig S1_1c) especially in recent years (2016-2017), we derived a new mean field in Fig S1_1d showing very small subtidal mean flow field within the bay.

With nine more drifter tracks obtained in 2018 and dozens proposed for 2019, we expect to get closer to defining the subtidal flow within the bay.

#### Forward tracks through modeled flow compared with drifter tracks

Surface drifter tracks from 2012 to 2015 are used as another metric to quantify the difference between FVCOM model currents and observed Lagrangian currents. We limited drifter data to those with sample intervals less than 2.5 hours. This resulted in 55 drifter tracks in Cape Cod Bay that had a total of 1365 drift days. We compared the model track to the drifter track by restarting the comparison every 3 days for a total of 454 comparisons. In other words, every three days, we restart the comparison at a drifter observation point. When the drifter trajectory is compared with the model trajectory, the "separation distance" and "distance ratio" are often used as two measures of accuracy. They are defined as the distance between the drifter point and the modeled track point at the same time and the ratio of separation distance to the drifter's total distance traveled, respectively. The results of observed vs modeled drifter tracks are shown in Fig S1_2. The 3-day "separation distances" are mainly distributed around 20 km, the minimum value is close to 0km and the maximum is 80 km or more; while the "distance ratios" are distributed around 0.2, the minimum value is close to 0 and the maximum value is above 1.0. Based on these comparisons, we estimate that the FVCOM model can be used to simulate particle pathways with an uncertainty of 200 meters per kilometer of trackline.

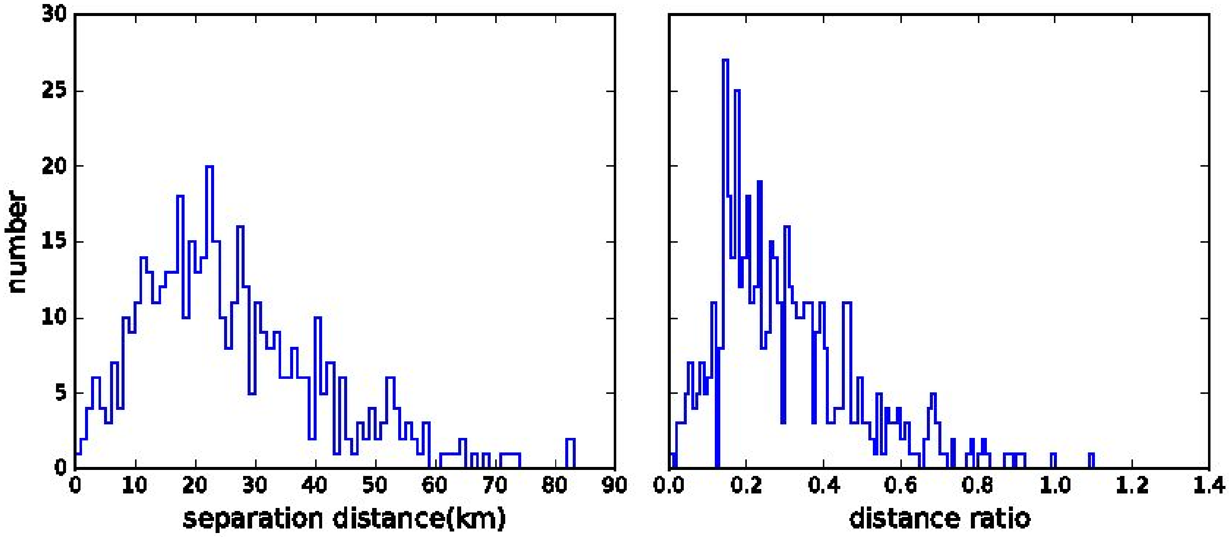


**Fig S1_2. The 3-day separation distance and the distance ratio between observed and modeled drifter track.**

#### Modeled vs Observed Temperatures

Taking 2012 and 2013 as an example, we compared the surface temperature observed at a

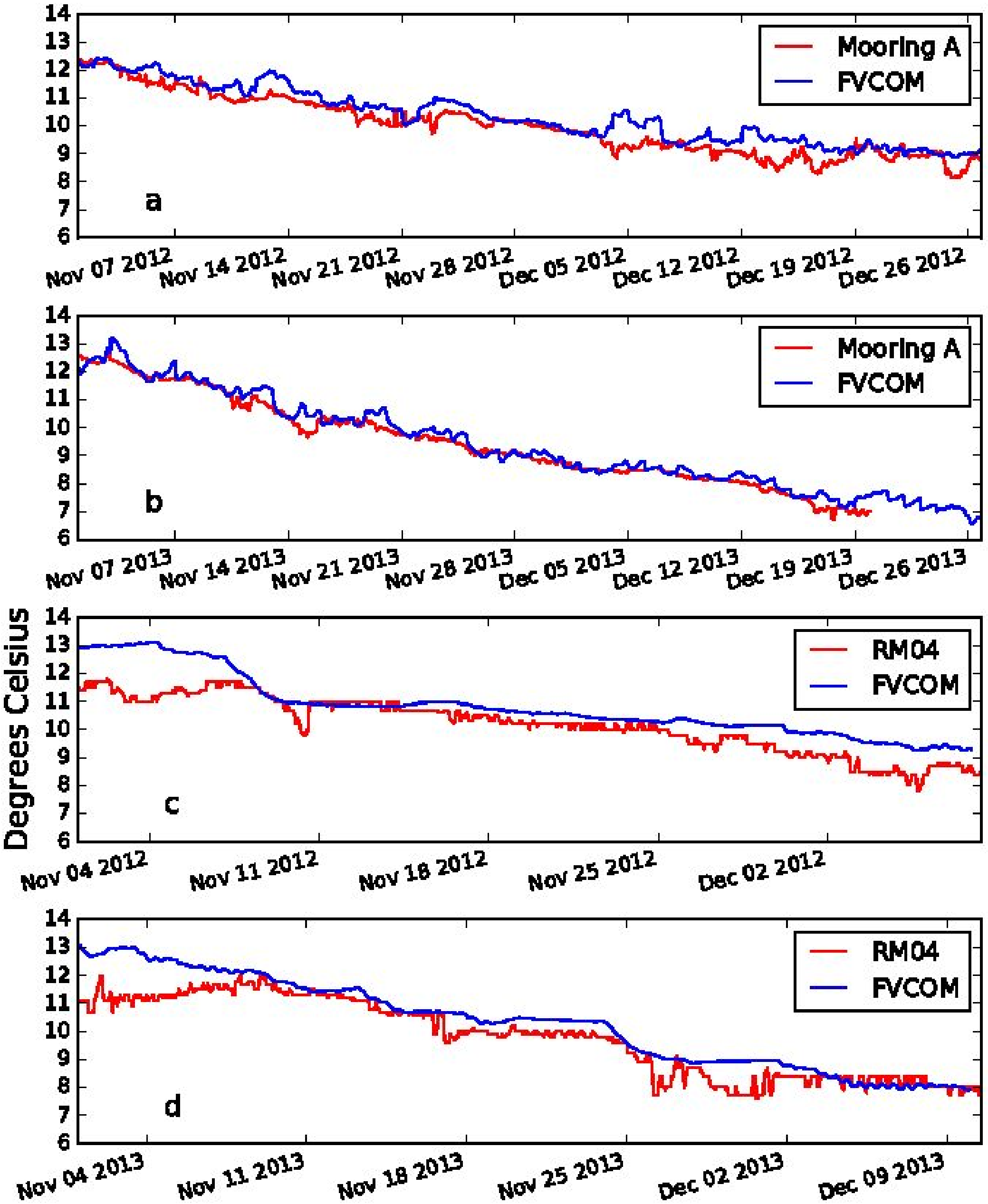


**Fig S1_3. Observed (NERACOOS Mooring A) vs modeled (FVCOM) surface (2 meters) temperatures in 2012 (a) and 2013 (b). Observed (eMOLT RM04) vs modelled (FVCOM) bottom temperatures in 2012 (c) and 2013 (d).**

mooring and the surface temperature simulated by the nearest FVCOM model grid cell (less than 1km away). As shown in Fig S1_3a and Fig S1_3b and Table S1_1, the surface temperature observed at NERACOOS mooring A is close (<0.4 ºC) to the temperature simulated by FVCOM.

We also compared the bottom temperature observed at a nearby (again <1km from model grid node) eMOLT station and the bottom temperature simulated by the FVCOM model, as shown in Fig S1_3c and Fig S1_3d. FVCOM bottom temperatures were less accurate near the bottom but still within 0.6 ºC of the observed. See Li et al (2017) for a more complete comparison of eMOLT bottom temperatures and FVCOM estimates.

Table S1_1. Observed minus modeled temperatures where NERACOOS "Mooring A" is observed at the surface and eMOLT "RM04" is observed at the bottom.

| Time | Index | Surface temperature | Bottom temperature |
|---|---|---|---|
| | | NERACOOS-FVCOM | eMOLT-FVCOM |
| 2012 | Mean | -0.385 | -0.540 |
| | Std | 0.334 | 0.517 |
| 2013 | Mean | -0.220 | -0.600 |
| | Std | 0.267 | 0.500 |
| 2014 | Mean | -0.087 | NA |
| | Std | 0.270 | NA |

In all of the above cases (2012-2014), it is clear that water temperature is important as well as the wind so it is necessary to understand the change of ocean temperature before estimating the origin of stranded turtles. Time series of surface temperature are extracted from 10 FVCOM grid nodes across Cape Cod Bay. The temperature across the 10 nodes is similar, dropping to 10.7 ºC on November 17  in both years. As is shown in this figure, there is very little spatial variability in the surface temperature across the bay and any one monitoring site could potential work as a marker of change.  If one site has cold stun conditions, they all do.

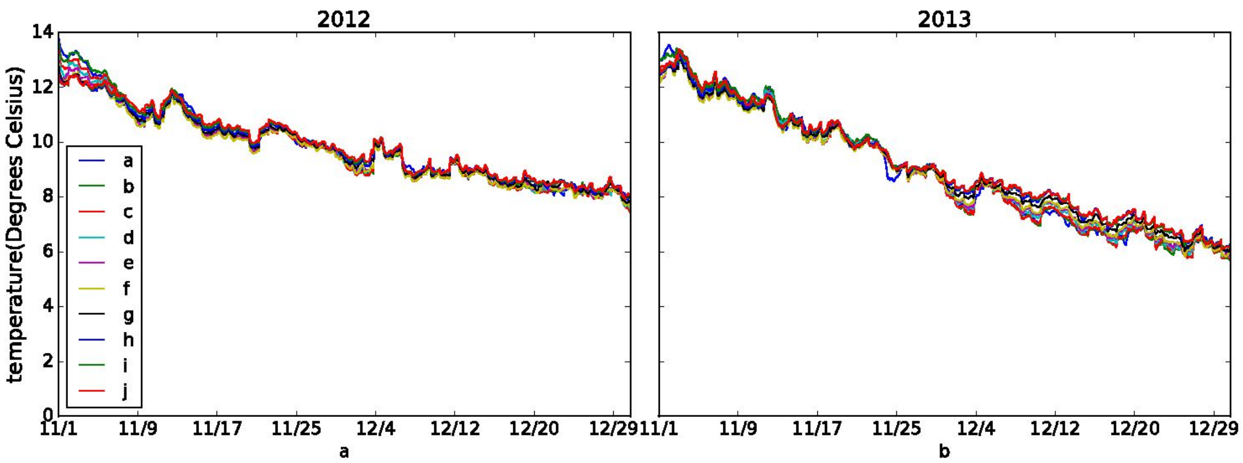


**Fig S1_4. Simulation of surface temperature for Cape Cod Bay and its adjacent waters in 2012 (a) and 2013 (b). The locations a-j are plotted in Fig 2 of the manuscript.**


Chen C, Liu H, and Beardsley RC. An unstructured, finite-volume, three-dimensional, primitive equation ocean model: application to coastal ocean and estuaries. J. Atm. & Oceanic Tech., 2003a;20: 159-186.

Li B, Tanaka KR, Chen Y, Brady D, and Thomas, AC. Assessing the quality of bottom water temperatures from the Finite-Volume Community Ocean Model (FVCOM) in the Northwest Atlantic Shelf region. Journal of Marine Systems 2017;173:21-30.
