## Supplementary material for "On simulating cold stunned turtle strandings on Cape Cod": https://www.nefsc.noaa.gov/epd/ocean/MainPage/turtle/ccbay/S2_Supplementary_material_CCBay_manuscript.pdf

### Appendix 2:

#### On methods to get turtles off the beach for particle tracking

As noted in the methods section, a variety of techniques were tested to get the stranded turtles off the beach in order to initiate backward particle tracking. Since most of the stranding reports have positions outside the model grid, we needed to define a new location some distance off the beach before the particle would move in the modelled flow fields. The code we used for this application is available at:

https://github.com/jamespatrickmanning/Lei_et_al_2019_CCBAY/tree/S2_method_off_beach.

The method we actually used is as follows and illustrated in Fig S2_1. The black line indicates the digitized coastline as defined by series of points. The spacing between these points is often different. The red point is the point where the turtle is stranded, the green point is the coastline point closest to the red point, and the blue line is the line between the stranded point and the nearest coast point. The yellow dot is a point where the green dot extends 1.5km along the blue line into the sea.

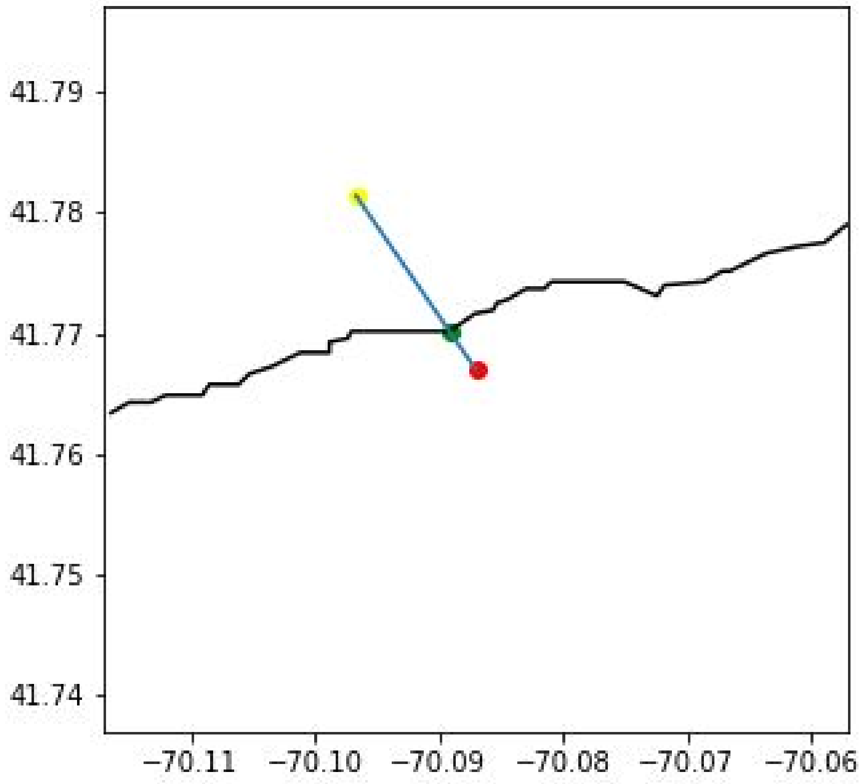


Fig S2_1. Simple method as used in the paper.

As shown in Fig S2_2 and S2_3, an alternative method was considered. This method generates a circle around the stranding (red point) with diameter “d=$X_1$ km” , calculates the two coastline points farthest from the red point in the circle, draws a straight line between them, calculates the midpoint of the line (green point), connects the red point and the green point, and extends $X_2$ km to the sea to get the yellow point. We found that the end result of this more complicated method was sensitive to a) the radius of the circle and b) the resolution of the coastline.

In Fig S2_2, $X_1$ = 1.7km and $X_2$ = 1.5km and in Fig. SM2_3, $X_1$ = 1.1km and $X_2$ = 1.5km. Due to the different radius of the circles, the yellow position points obtained in Fig SM2_2 and Fig S2_3 are significantly different. In cases of very complex coastline, neither method will provide ideal results for each stranding. As noted in the manuscript, some tracks were abandoned when they failed to advect offshore using the simpler method.

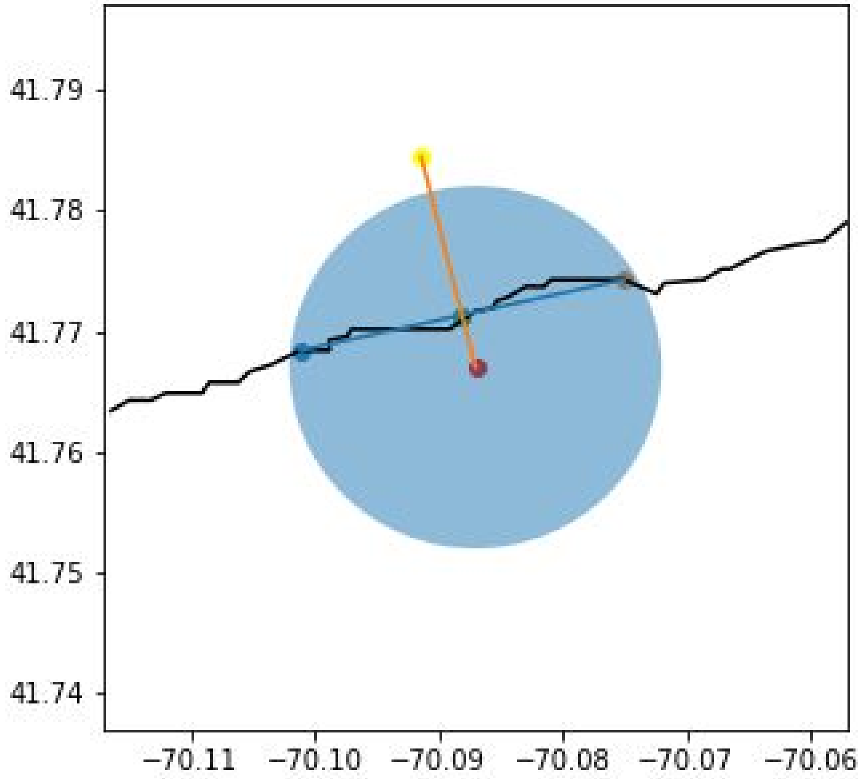


Fig S2_2. Complex method (The radius of the circle is 1.7km)

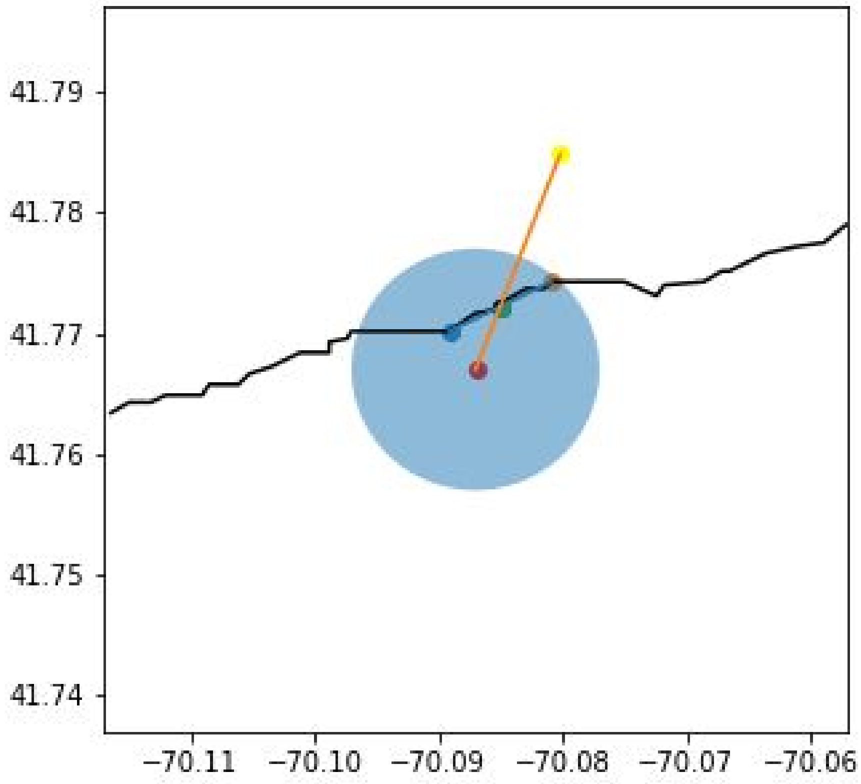


Fig S2_2. Complex method (The radius of the circle is 1.1km)
