## Supplementary material for "On simulating cold stunned turtle strandings on Cape Cod": https://www.nefsc.noaa.gov/epd/ocean/MainPage/turtle/ccbay/S3_Supplementary_material_CCBay_manuscript.pdf

### Appendix 3: On animating drifter tracks and wind with Python

In order to best visualize the variability of circulation in Cape Cod Bay, we animated the drifter tracks and overlaid a representation of wind in the form of a web-served gif file. The code for this animation can be found here. It uses two input files, the ascii data for a particular batch of drifters tracks and a time series of NCEP wind for a nearby grid point. The drifter data can be found at NOAA's Northeast Fisheries Science Center page here and the NCEP wind was downloaded from their FTP site. After running the Python code "animate_drifter_Basemap.py" we have a set of ordered frame*.png files in a particular directory where we then run the Linux "convert -delay 10 -loop 0 frame*.png /net/pubweb_html/drifter/drift_audubon_2018_1.gif", for example. The resulting animation can be found at:

https://www.nefsc.noaa.gov/drifter/drift_audubon_2018_1.gif
